# Maternal IL-17A administration fails to disrupt fetal cortical lamination in the absence of maternal microbes

**DOI:** 10.64898/2025.12.15.693075

**Authors:** Izan Chalen, Selena Wang, Rafael J. Gonzalez-Ricon, Ashley M. Otero, Fernando J. Rigal, Alexander Byrne, Shakirat Adetunji, Catherine Best-Popescu, Adrienne M. Antonson

**Affiliations:** Department of Animal Sciences, University of Illinois Urbana-Champaign, Urbana, IL, USA; Indiana University School of Medicine, Indianapolis, IN, USA; Neuroscience Program, University of Illinois Urbana-Champaign, Urbana, IL, USA; Department of Physics, University of Illinois Urbana-Champaign, Urbana, IL, USA; School of Molecular and Cellular Biology, University of Illinois Urbana-Champaign, Urbana, IL, USA; College of Veterinary Medicine, University of Illinois Urbana-Champaign, Urbana, IL, USA; Department of Bioengineering, University of Illinois Urbana-Champaign, Urbana, IL, USA

**Keywords:** Germ free, cortical lamination, fetal neurodevelopment, IL-17A, placenta, maternal immune activation, maternal microbiome

## Abstract

Maternal inflammation during pregnancy can disrupt fetal neurodevelopment and contribute to the pathogenesis of neurodevelopmental disorders. Interleukin-17A (IL-17A) has emerged as a potential mediator linking maternal immune activation (MIA) to neurodevelopmental abnormalities in offspring. Production of this cytokine is in-part regulated by the maternal gastrointestinal microbiome. However, it remains unclear whether IL-17A alone is sufficient to induce offspring brain abnormalities independent of maternal microbial signals. This study tests the potential for IL-17A to mediate fetal brain cortical architecture through administration of recombinant (r)IL-17A to pregnant germ-free (GF) dams. Pregnant GF mice received daily intraperitoneal injections of rIL-17A for six days, covering the mid- to latter half of the gestational period. Placental tissue was processed for histopathological evaluation, and cortical lamination was examined in the fetal brain using gold-standard immunohistochemical approaches. Spatial Light Interference Microscopy (SLIM), a novel label-free imaging technique, was employed to further quantify cortical architecture. This is the first report of SLIM imaging performed in the mouse embryonic brain. Overall, administration of rIL-17A to pregnant GF dams during mid-to-late gestation resulted in normal placental morphology and fetal cortical lamination patterns, suggesting that IL-17A alone is insufficient for altering cortical neurodevelopment, at least at this gestational timepoint. These findings highlight the importance of accounting for the maternal microbiome when interpreting the impacts of prenatal inflammation on brain development *in utero*.

## 1.0 Introduction

The prenatal period is a sensitive window in which developmental processes determine health outcomes in later stages of life [1–3]. In the 1980s, the epidemiologist David Barker first showed that low birth weight—due to poor fetal growth or nutrition—led to increased risk of later life cardiovascular disease; he proposed the Fetal Origin of Adult Disease hypothesis, subsequently known as the Barker Hypothesis [1–4]. Further evidence supporting Barker’s hypothesis has shown that neurodevelopmental disorders (NDDs) occur at higher rates following early life insults [3, 5–11]. Notably, environmental insults that disrupt the early immune milieu are consistently linked to NDDs, especially during windows of plasticity, when organisms exhibit heightened sensitivity to immune perturbations [5, 12, 13]. Among these immune perturbations, exposure to maternal infection during gestation is one of the strongest risk factors predisposing individuals to NDDs [14, 15]. To investigate the cellular and molecular mechanisms underlying this risk, researchers frequently employ the maternal immune activation (MIA) model, experimentally inducing inflammation during pregnancy to examine downstream signaling patterns that impact offspring brain and behavioral outcomes [6, 12, 13, 16–20]. Many of these studies have revealed shared patterns of cytokine signaling independent of the immune stimulant, leading to the hypothesis that an imbalance in a select few cytokines could be driving adverse neurodevelopmental outcomes [21]. Several recent studies support the possibility that interleukin (IL)-17A could be one of these major drivers, altering cortical patterning and ultimately disrupting the neuronal circuitry that governs NDD-like behaviors [22–27]. Beyond pathogens (or pathogen mimetics like synthetic viral RNA analog poly I:C or purified bacterial lipopolysaccharides), combined environmental stressors such as air pollution and maternal stress also activate the maternal immune system and elevate IL-17A, leading to long-lasting, male-specific impairments in social behavior and microglial function [13, 28].

Once in the brain, IL-17A has been shown to play a broad role in neurodevelopmental and neuroimmune regulation via binding receptors on both neurons and glia [29–31]. Neurons express the IL-17RA/RC receptor complex, and fetal brain *Il17ra* transcripts have been shown to be upregulated following maternal poly I:C exposure, indicating that neurons are capable of sensing this cytokine during corticogenesis [22, 31]. Engagement of these receptors has also been shown to alter neuronal signaling, leading to subsequent synaptic dysfunction and blood–brain barrier breakdown [31, 32]. In addition, IL-17A acts directly on microglia, perpetuating further release of inflammatory cytokines and reactive oxygen species, amplifying neuroinflammatory cascades [29, 30]. Taken together, these findings highlight why IL-17A exposure during pregnancy may be particularly consequential for offspring neurodevelopmental trajectories.

Emerging evidence suggests that another critically important factor influencing prenatal development is the maternal microbiome. The fetus requires signals from the maternal microbiota to support proper placental and neurological development [33–37]. These maternal microbial-derived signals include specific metabolites—such as trimethylamine N-oxide (TMAO), 5-aminovalerate (5-AV), indole-3-propionate (IP), and hippurate (HIP)—which have been shown to promote axon outgrowth in fetal thalamic explants and rescue axonogenesis *in vivo* [34, 38]. During MIA, the maternal microbiome—particularly segmented filamentous bacteria (SFB)—has been shown to be required for engaging intestinal T helper (T_H_)17 cells following a viral mimic poly I:C challenge [22, 23, 37, 39–41]. These maternal T_H_17 cells are the primary producers of IL-17A in the poly I:C-induced MIA model, and IL-17A appears to directly disrupt the developing somatosensory cortex. Blocking IL-17A or genetically knocking down T_H_17 cells during MIA both protect the offspring from cortical malformations and NDD-like behaviors [22]. Notably, the characteristic fetal cortical abnormalities described in these studies were dependent on the presence of SFB [12, 42, 43]. An SFB-rich microbiota increased circulating IL-17A following poly I:C challenge, resulting in classic NDD-like behaviors in offspring. Conversely, mice who lacked SFB in their microbiome failed to mount this IL-17A response, and their offspring failed to develop NDD-like behaviors [43].

Interestingly, MIA has also been shown to alter the gastrointestinal microbiota of offspring, including reduced microbial diversity and taxa such as *Clostridia* and *Bacteroides* enrichment [44–46]. These shifts resemble microbial features reported in human patients with autism spectrum disorder, where decreased diversity and overrepresentation of certain anaerobic bacteria are recurrent findings [44–46]. Furthermore, maternal age-associated gut dysbiosis can create long-lasting behavioral abnormalities in the offspring, concomitant with shifted fecal microbial profiles and altered metabolite and neurotransmitter signatures [47]. Taken together, this evidence shows the importance of the crosstalk between the central nervous system, the microbiome, and the immune system during development [22, 37, 39–41].

The multi-layered and highly vascularized placenta constitutes the maternal-fetal interface, serving as a key regulator of fetal development through modulating selective passage of maternal signals into the fetal compartment [48–51]. Importantly, its structural integrity and innate immune defenses help prevent vertical transmission of pathogens to the developing fetus [52–55]. Still, maternal inflammation can alter placental morphology and physiology, with potential negative consequences for fetal development, including atypical neurodevelopment [52, 53, 56–60]. This effect is partly shaped by the maternal microbiome, which promotes normal placental growth and vascularization [35].

Building on prior evidence that microbial signals shape both placental and neural development, this study aims to test whether IL-17A, administered to (germ-free) GF pregnant dams in the latter half of gestation, is sufficient to disrupt fetal cortical development in the absence of microbes. We use GF mice to disentangle any microbial-derived signals from IL-17A-specific signaling. Using a novel label-free imaging technique called Spatial Light Interference Microscopy, we complement traditional immunohistochemistry measures of cortical brain structure to assess lamination patterns in late-gestation fetuses. Additionally, we analyze placenta morphology and pathology to assess the integrity of the maternal–fetal interface. Collectively, our findings show that repeated maternal IL-17A administration is insufficient to disrupt fetal cortical lamination in the absence of microbial signals.

## 2.0 Methods

### 2.1 Animals

All animal work was performed at the University of Illinois Urbana-Champaign’s Rodent Gnotobiotic Facility. UIUC Animal Care Technicians routinely test samples from in-house germ-free mice via microbial culture to confirm GF status. 23 nulliparous 8-to-10-week-old female C57BL/6 GF mice were bred in pairs to GF male studs across two biological replicates. The presence of a vaginal plug was designated as gestational day (GD) 0.5. Recombinant IL-17A (rIL-17A; R&D Systems, Catalog no: 7956-ML-010/CF) or an equivalent volume of PBS was administered intraperitoneally (i.p.) to pregnant dams once per day from GD10.5 to 15.5 between the hours of 7:00 and 9:00 AM. A dose of 10 µg/mL of rIL-17A in 100µL PBS (1 µg/mouse) was selected based on previous publications investigating intestinal inflammation [61–63]. We targeted this mid-to-late gestational window as it aligns with the developmental timepoint when MIA is known to cause aberrant fetal cortical phenotypes [22, 23, 34]. At GD16.5 (24 h after the last injection), fetal brains and placentae were collected and fixed.

To establish IL-17A concentrations in circulation following intraperitoneal delivery, nine male GF mice (13-14 weeks old) were injected i.p. with 1 µg of rIL-17A (as above). Blood was collected at predetermined time points: 0, 15, 30 min, and 1, 2, 4, 6, 12, and 24 h post-injection. Serum levels of IL-17A were measured to determine a time course.

All animal experiments were performed in the Rodent Gnotobiotic Mouse Facility at the University of Illinois Urbana-Champaign. Animals were maintained on a 12 h light–dark cycle. All animal research was approved by and performed in accordance with the Institutional Animal Care and Use Committee (IACUC) at the University of Illinois Urbana-Champaign.

### 2.2 Tissue collection

Tissues were collected under sterile conditions at GD 16.5. Pregnant dams were euthanized by CO_2_ asphyxiation. Maternal blood was obtained through cardiac puncture. The uterus was excised and placed in cold PBS, then placentae and fetuses were separated and immersed whole in 10% neutral buffered formalin (NBF) and stored at 4 °C. Maternal spleen mass was obtained, and colon length was measured after the full gastrointestinal tract was excised. Whole blood was centrifuged at 2000 × *g* at 4 °C for ten minutes, and the serum was stored at –80 °C until analysis.

### 2.3 Measuring serum IL-17A concentrations

IL-17A concentrations in serum were measured using an ELISA MAX™ Deluxe Set Mouse IL-17A kit from Biolegend (Catalog no: 432504) following the manufacturer’s protocol.

### 2.4 Placental histopathological scoring

Fixed placentae were washed in PBS, followed by paraffin-embedding and transverse sectioning at 5 µm using a microtome. The tissue was mounted on slides and allowed to dry on a slide heater at 55°C. Afterwards, the tissue was stained with Hematoxylin and Eosin (H&E) as follows: deparaffinization and rehydration with gradients of xylenes and ethanol (2 min in each), nuclear staining with hematoxylin (2 min), differentiation in acid alcohol, bluing in ammonia water (30 sec), cytoplasmic counterstain with eosin (30 sec), followed by dehydration, clearing, and coverslip mounting with Fluoromount-G mounting medium (Fisher Scientific, Catalog no. 50 187 88) [64]. For each sample, three placenta sections were selected from the centermost region and imaged under the bright-field function of a Keyence BZ-X810 microscope. The images were stitched using ImageJ software using the stitching plugin [65]. Using ZEISS ZEN 3.0 software (Oberkochen, DE), the area of each of the regions of interest (labyrinth, junctional zone, and decidua) in the placenta was measured by manually tracing each region. Once that area was obtained, the ratio was calculated by dividing the area of the labyrinth by the area of the junctional zone; this indicator has been previously reported as a proxy for placental efficiency [48–50].

Additionally, placentae were scored for necrosis, mineralization, and leukocyte infiltration by board-certified pathologist Dr. Shakirat Adetunji, who was blinded to treatment group. H&E-stained slides were examined on a brightfield microscope (Olympus BX45TF), and pathology was assessed using a semi-quantitative rubric adapted from established diagnostic criteria [53, 66]. Scored features included: mineralization, necrosis, mononuclear leukocytes, and segmented leukocytes. For mineralization, grades were defined as 0 = none or rare scattered small foci; 1 = mild, multifocal; 2 = moderate, multifocal to coalescing within the decidua and/or extending into additional placental zones; 3 = large multifocal or diffuse mineralization within a zone. For necrosis, grades were defined as 0 = none or rare scattered small foci; 1 = mild, multifocal confined to the decidua; 2 = moderate, multifocal in the decidua and/or other zones; 3 = multifocal to coalescing, large necrotic areas in the decidua and/or other zones. For inflammatory cells, mononuclear leukocytes and segmented leukocytes were scored separately using: 0 = none/normal; 1 = slight increase; 2 = moderate increase; 3 = marked increase. Lesion distribution was noted across decidua basalis, junctional zone, and labyrinth. Each feature was graded on a 0–3 scale with the following anchors: 0 = none/rare; 1 = mild, multifocal; 2 = moderate, multifocal to coalescing and/or involving additional placental zones; 3 = large multifocal or diffuse involvement within a zone. Three non-serial sections (50 µm apart) from one representative placenta per litter were evaluated, and feature scores were averaged across sections to obtain a per-litter mean used for statistical analyses.

### 2.5 Fluorescent immunohistochemistry in fetal brains

Fixed fetuses were washed in PBS, followed by excision of the fixed fetal brain from the skull. Fetal brains were immersed in 30% sucrose solution at 4°C until the tissue sank. Cryoprotected fetal brains were embedded in OCT and cryosectioned to obtain 50 µm coronal sections. Free-floating batch staining was performed following a previously described protocol [67]. Sections were washed three times in PBS with 0.05% Tween-20 (Thermo Fisher Scientific, Catalog no. PRH5152; PBST) for 5 min each time and subsequently incubated with blocking buffer (5% goat serum [R&D Systems, Minneapolis, MN, Catalog no. S13110], 1% bovine serum albumin [Thermo Fisher Scientific, Catalog no. 126609100GM], 0.3% Triton-X 100 [Thermo Fisher Scientific, Catalog no. ICN19485450] in PBST) for 1 h at room temperature. Sections were incubated overnight at 4 °C with primary antibodies: rabbit anti-TBR1 (1:1000; Abcam, Cambridge, UK, Catalog no. ab183032), or rabbit anti-IL17RA (1:1000; Thermo Fisher, Catalog no. PA5-34571). Then, sections were washed three times in PBST and incubated with secondary antibodies: Alexa Fluor 594 goat anti-rabbit IgG H&L (1:250; Jackson ImmunoResearch, West Grove, PA, Catalog no. 111-585-003) for 2 h followed by staining in DAPI (Thermo Fisher Scientific, Catalog no. EN62248) for 1 min. Sections were mounted with Fluoromount-G Mounting Medium (Thermo Fisher Scientific, Catalog no. 5018788) and stored at 4 °C until imaging.

### 2.6 Quantification of fluorescent immunohistochemistry

Sections were imaged on a ZEISS AxioScan.Z1 slide scanner using a 20X/0.8 NA objective; image files were blinded and analyzed in ZEN 3.0 Blue software, where mean fluorescence intensity (MFI) of each marker (TBR1 or IL-17RA) was quantified within a region of interest (ROI) in the fetal neocortex. For DAPI quantification, mean fluorescence intensity was measured in ZEN 3.0 Blue within the same ROI, subdivided into cortical layers I, II/III, IV/V, and VI, to estimate nuclear density across laminae. The ROI was a 300 × 300 μm² square in the cortical plate encompassing the dysgranular zone of the primary somatosensory cortex (S1DZ). This region was selected based on previously published studies by our group and others [22, 67]. The ROI was registered to a fixed position by distance from the midline and the retrosplenial cortex relative to the dorsal midline length so that the same anatomical position was imaged across sections, following methodology from previous publications [22, 67]. Thickness of the cortical plate was measured as previously described [67] using ZEISS ZEN 3.0 Blue. For laminar profiling of TBR1, the ROI was divided into 10 equal bins as in previous publications [22, 67]. For subsequent counts of individual TBR1+ neurons, ROI images were exported, and events were quantified in Biodock.ai [68], described in detail below.

### 2.7 Spatial Light Interference Microscopy (SLIM)

A subset of fixed fetal brains were embedded in paraffin and sectioned at 5 µm using a microtome. Sections were mounted on slides and stained with H&E using a brief, standardized sequence, as described in detail above: deparaffinization and rehydration, nuclear staining with hematoxylin, differentiation in acid alcohol, bluing in ammonia water, cytoplasmic counterstain with eosin, followed by dehydration and cover slipping [64].

Using a Leica DMi8 microscope with a SLIM module, quantitative phase images were acquired from four sequential intensity frames with π/2 phase shifts (0, π/2, π, 3π/2) under white-light illumination at 40X magnification, and tiled fields (∼8% overlap) were stitched to generate wide-field maps [69]. Images of the ROI were obtained under a 10X magnification, then processed using Biodock (available from www.biodock.ai) [68] for event quantification. The ROI was 300 × 300 µm² square in the cortical plate representing the somatosensory cortex, as described above; the same anatomical position was imaged across sections.

### 2.8 Quantification of IHC, and SLIM images using Biodock.ai

Images from traditional IHC and label-free SLIM, were processed using the online artificial intelligence software Biodock.ai [68]. We uploaded the ROI images to the system and created a project for automatic counting. The ROI was subdivided into 1000-pixel-wide square tiles. The first step of the project was to train the system by manually selecting every identifiable “event” (i.e., cell nucleus or body) in a tile, then the system would replicate the counting in another tile, and then manual corrections were made. Once the system was trained, all the ROI images were analyzed by the system (sufficient training was verified manually by an experienced scientist). The threshold function was used to distinguish individual events within each section, defining groups of morphologically similar structures based solely on their shape and intensity characteristics. To validate this approach, we manually reviewed system-generated classifications and corrected mis-assignments until the output matched the ground-truth cell counts across representative tiles. Analysis of fluorescent IHC, and SLIM images was performed independently.

### 2.9 *In vitro* test of rIL-17A bioactivity

Preserved BV-2 microglial cells (derived from C57BL/6 mice, donated by Dr. Robert McCusker) were thawed at 37 °C for 1–2 min, transferred under aseptic conditions to a 15 mL conical tube containing 9 mL of pre-warmed complete culture media with the following composition: DMEM (Dulbecco’s Modified Eagle Medium; Corning, catalog number: 10-013-CM) supplemented with 10% (v/v) FBS (Fetal Bovine Serum; Corning, catalog number: MT35010CV), 1% (v/v) Pen/Strep (ThermoFisher, catalog no: 15140122); and centrifuged at 300 × g for 3 min. The pellet was resuspended in fresh culture media and cells were counted using trypan blue exclusion. Cells were seeded at a density of 1×10⁶ per well in 6-well plates with 2 mL of complete media and allowed to adhere overnight at 37 °C with 5% CO₂. A total of two plates were prepared to accommodate two technical replicates. Each plate included the following conditions: (*1*) negative control (PBS-treated cells), (*2*) positive control (1 µg/mL LPS), (*3*) low-dose rIL-17A (1 ng/mL), and (*4*) high-dose rIL-17A (100 ng/mL). The low- and high-dose rIL-17A concentrations were selected based on previous publications [61–63] and on our own time-course data **(see Fig. 1I)**. Treatments were prepared by diluting rIL-17A or lipopolysaccharides (LPS from *E. coli* O55:B5, Millipore Sigma, catalog number: L288-10MG) into pre-warmed culture media immediately prior to use. At 12h post-stimulation, supernatants were collected from each well and stored at –20 °C until analysis.

**Figure 1.**
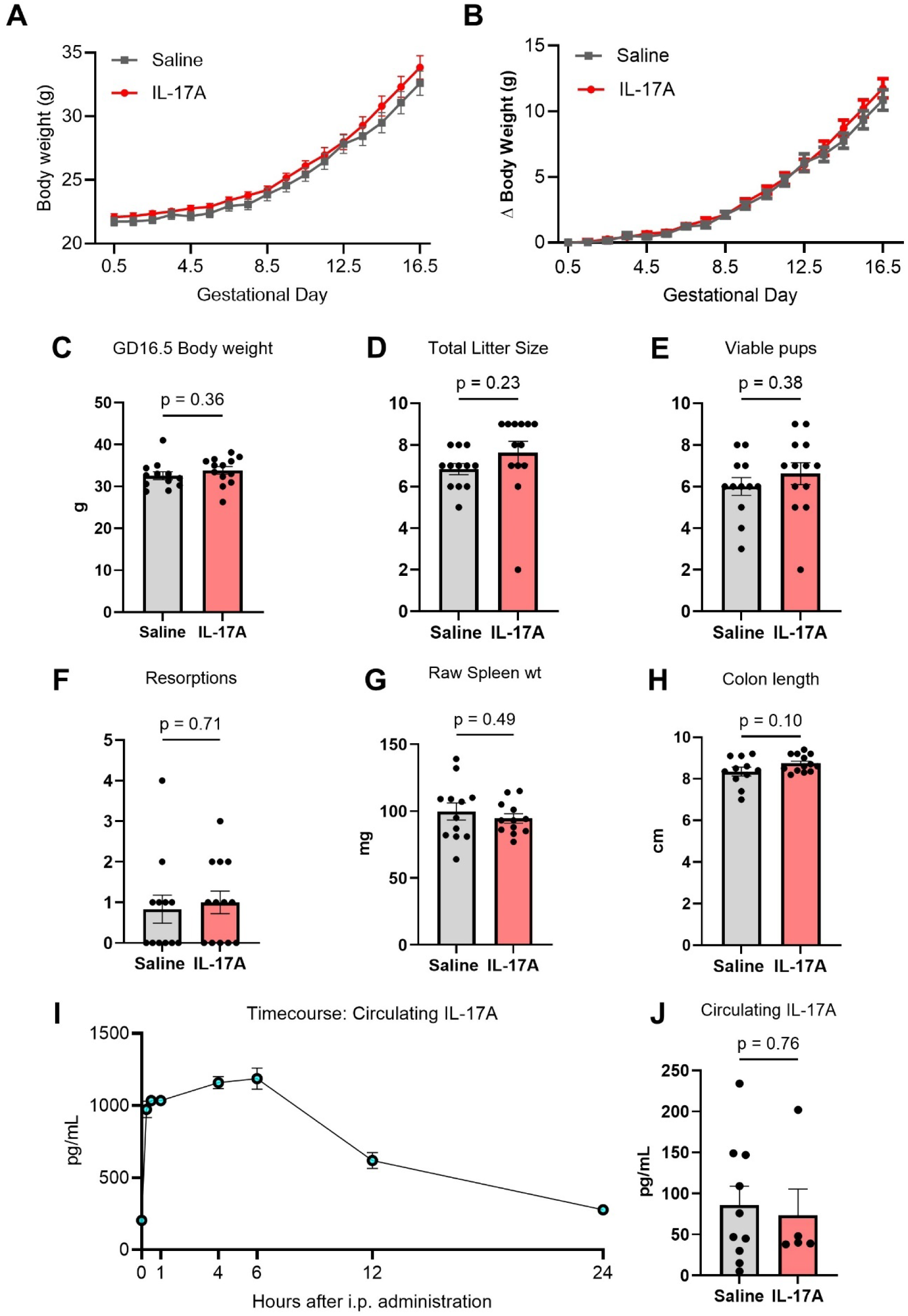
Gestational IL-17A administration did not alter pregnancy outcomes. Administration of 1 µg recombinant IL-17A i.p. daily (GD10.5–15.5) did not affect **(A)** maternal body weight trajectory nor **(B)** total body weight gain during pregnancy. **(C)** Maternal body weight at GD16.5 was unchanged by IL-17A treatment. **(D)** IL-17A administration did not affect total litter size. **(E)** IL-17A administration did not affect the number of viable pups per litter. **(F)** IL-17A administration did not affect the number of resorptions per litter. **(G)** Maternal spleen weight was not significantly altered by IL-17A treatment. **(H)** Maternal colon length was not significantly altered by IL-17A treatment. **(I)** Time-course analysis of circulating IL-17A after a single intraperitoneal injection confirmed systemic exposure, peaking within hours and returning to baseline by 24 h. **(J)** At GD16.5, 24 h after the final injection, circulating IL-17A levels did not differ between saline- and IL-17A-treated dams (Saline: n = 10; IL-17A: n = 5). Data are means ± SEM. Saline: n = 12-13; IL-17A: n = 12 (unless otherwise stated).

### 2.10 Quantification of protein in cell supernatants

Supernatant from *in vitro* BV-2 experiments was assayed using the Proteome Profiler Mouse Cytokine Array (R&D Systems, catalog no: ARY006). Following the kit instructions, 2 mL of the block buffer was pipetted into each well of the 4-well multi-dish, then each membrane was placed in a separate well facing upwards. After one hour of incubation on a rocking platform shaker, the block buffer was removed and replaced by a 1.5 mL mixture containing 1 mL of sample (pooled supernatant from each experimental condition) and 0.5 mL of array buffer. After overnight incubation at 4 °C the membranes were individually washed three times in 1X wash buffer for ten minutes on a rocking shaker. Then diluted antibody cocktail was added to the membranes. After incubation for one hour at room temperature, the membranes were washed again three times and excess wash buffer removed. Then the membranes were incubated at room temperature for one hour in Streptavidin-HRP. After washing the membranes three times again, the membranes were placed on the bottom sheet of a plastic sheet protector and covered with 1 mL of Chemi Reagent Mix, then incubated for one minute while covered by the plastic sheet protector. The membranes were then developed for three minutes on an iBright1500 system (Invitrogen). The pixel density of each target was measured using ZEISS ZEN 3.0 Blue software.

### 2.11 Statistics

Dam was treated as the experimental unit for all outcomes, and a minimum of one randomly selected fetus per litter was used for each experimental outcome. All data were analyzed using GraphPad Prism 9 Software (San Diego, CA), with significance set at α = 0.05. Treatment groups were compared using unpaired t-tests, non-parametric Mann-Whitney U tests (for discrete data) or repeated measures ANOVA, as appropriate. Welch’s t-test was used when variances were unequal. All data are expressed as means ± SEM.

## 3.0 Results

### 3.1 Administration of IL-17A does not alter maternal health nor litter characteristics

To assess the effects of rIL-17A administration on pregnant GF dams, body weight was recorded daily. We did not observe a difference in body weight across pregnancy **(Fig. 1A)** nor in daily weight gain **(Fig. 1B)**. At the end of the experiment, the weight of dams in both groups was not different **(Fig. 1C)**. Similarly, administration of rIL-17A did not impact total litter size **(Fig. 1D)** nor total number of viable pups **(Fig. 1E)** or resorptions per litter **(Fig. 1F)**.

Maternal spleen weight, which is commonly used as an indicator of systemic inflammation [70, 71], did not differ between controls and IL-17A groups **(Fig. 1G)**. Maternal colon length, a common proxy for intestinal inflammation or colitis [72], similarly did not differ **(Fig. 1H)**.

To measure circulating concentrations of IL-17A post-injection, we performed an independent time-course analysis in a subset of GF animals. This experiment confirmed that rIL-17A administered i.p. enters circulation within 15 min, with serum levels peaking at 6 h (mean of 1191 ± 33 pg/mL) and returning to near-baseline levels (mean of 277 ± 24 pg/mL) within 24 h **(Fig. 1I)**. Notably, the peak circulating IL-17A concentration is in line with serum levels reported in poly I:C-induced MIA [22, 39]. As we did not want to add the additional confounding stress of repeated blood collection from pregnant experimental animals, circulating levels of IL-17A in experimental dams were only measured at the time of tissue collection, 24 h after the final injection (GD16.5). Consistent with the return to baseline observed at 24 h in the time course experiment **(Fig. 1I)**, we did not observe any differences in circulating levels of IL-17A in pregnant dams at GD16.5 **(Fig. 1J)**. This indicates that rIL-17A was likely cleared from circulation each day, and that repeated daily injections did not result in systemic accumulation of the cytokine over time.

To confirm the biological activity of rIL-17A, we measured IL-17A–stimulated responses *in vitro* using murine BV-2 cells. This microglia-like cell line served as a proxy for measuring the ability of fetal brain cells to respond to local rIL-17A. Doses of rIL-17A were selected based on our *in vivo* time-course data, whereby the low dose (1 ng/mL) reflected the physiological peak in circulation (Fig. 1I), and the high dose reflected a supraphysiological concentration to test the upper limit of responsiveness [73]. As an internal control, responsiveness to bacterial endotoxin LPS (1 µg/mL) stimulation was measured. Both IL-17A doses modulated the BV-2 response, affecting ICAM-1, IL-13, CCL3, CCL4, TNF-α, IL-1ra, CCL12, and CXCL10 production **(Supp. Fig. S1A)**. Levels of IL-17 recovered from supernatant reflected the low and high doses administered across conditions **(Supp. Fig. S1C)**, and LPS initiated the expected innate anti-bacterial pro-inflammatory response **(Supp. Fig. S1C)**. Together, these data confirm bioactivity of the rIL-17A administered *in vivo*.

### 3.2 Administration of IL-17A does not overtly impact GF placental histopathology or zonal structure

As maternal immune status may alter placental physiology, and subsequent fetal development, we assessed placental health in GF dams in response to IL-17A using standard histological examination practices. First, we measured the area of each distinct tissue layer of H&E-stained placentae: decidua, junctional zone, and labyrinth **(Fig. 2A,B)**. Overall, placental area did not statistically differ between the experimental groups, although average area was numerically smaller following IL-17A administration (p = 0.077; Fig. 2C). Similarly, the junctional zone **(Fig. 2D)** and labyrinth areas **(Fig. 2E)** were numerically smaller on average following IL-17A compared to controls, although this did not reach significance (p = 0.075 and p = 0.11 respectively).

**Figure 2.**
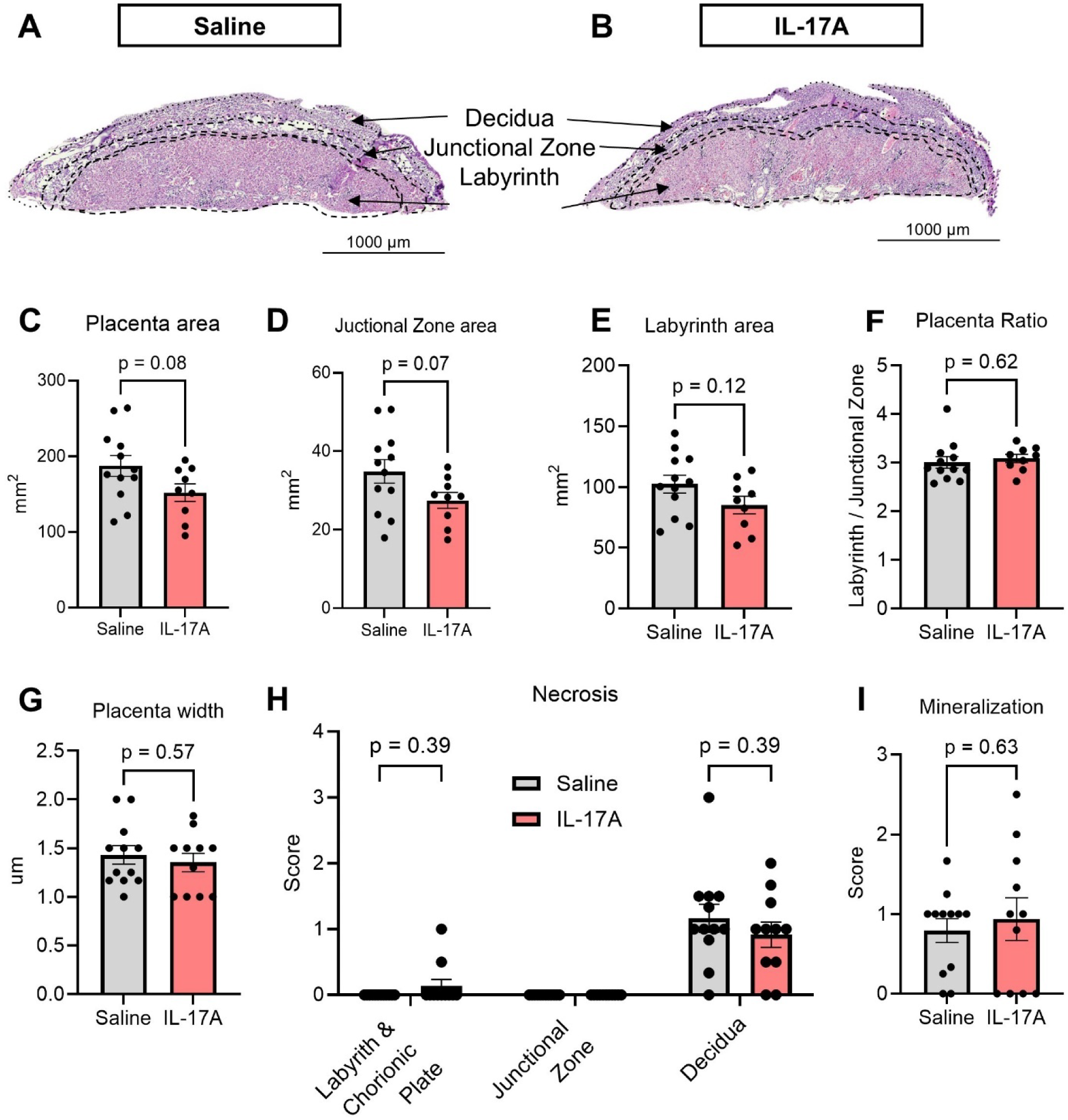
Gestational IL-17A administration did not alter placental morphology nor histopathology. Representative H&E-stained transverse sections of mouse placentae from a **(A)** saline-treated dam and **(B)** IL-17A-treated dam, showing the decidua, junctional zone, and labyrinth. Scale bar = 1000 µm. **(C)** IL-17A administration did not affect the total placental area in medial sections. **(D)** IL-17A administration did not affect the junctional zone area. **(E)** IL-17A administration did not affect the labyrinth area. **(F)** IL-17A administration did not affect the labyrinth-to-junctional zone area ratio. **(G)** Placental width measured at the medial section did not differ between groups. **(H)** Semi-quantitative scoring of necrosis across the labyrinth, chorionic plate, junctional zone, and decidua revealed no differences between groups. **(I)** Placental mineralization was not altered by IL-17A administration. Data are means ± SEM. Saline: n = 9-12; IL-17A: n = 11-12.

We used the ratio of the labyrinth to the junctional zone as a morphological proxy for feto-maternal vascular exchange capacity, whereby a relatively larger labyrinth indicates greater surface area for nutrient and gas transfer and thus higher placental efficiency. Disruptions in this ratio have been reported in various conditions of placental insufficiency [48–50]. However, there was no difference in this ratio between control and IL-17A groups **(Fig. 2F)**, nor did overall placental width differ **(Fig. 2G)**, indicating that overt placental structural abnormalities are absent in this context.

To examine more specific indicators of placental disruption as a consequence of IL-17A administration, placentae were scored for inflammatory lesions by a board-certified pathologist. Consistent with the structural observations, there were no differences in necrosis across any of the three regions analyzed (labyrinth and chorionic plate, junctional zone, and decidua; **Fig. 2H**). Similarly, no notable differences were observed in mineralization **(Fig. 2I)**, an indicator of malperfusion, and placentae in both groups lacked segmented and mononuclear leukocytes (data not shown as scores were all zero). Taken together, these histological findings indicate that IL-17A administration in GF dams did not shift placental physiology.

### 3.3 Administration of IL-17A in GF pregnant mice does not alter the structure of the fetal somatosensory cortex

In order to assess the structural integrity of the developing somatosensory cortex, we measured the patterning and abundance of TBR1+ deep-layer glutamatergic neurons across ten equally distributed bins corresponding to descending cortical depths **(Fig. 3A)**. Absence or disorganization of TBR1 in the somatosensory cortex has been reported in poly I:C-induced MIA [22, 23]. Here, there was no difference in TBR1⁺ cell distribution across any of the ten bins as quantified by individual cell number **(Fig. 3B)**. Normalized mean fluorescence intensity (MFI) was also unchanged **(Fig. 3C)**. Within the whole ROI, neither total TBR1+ cells **(Fig. 3D)** nor MFI **(Fig. 3E)** differed. The overall thickness of the cortical plate also did not differ between control and experimental groups **(Fig. 3F)**.

**Figure 3.**
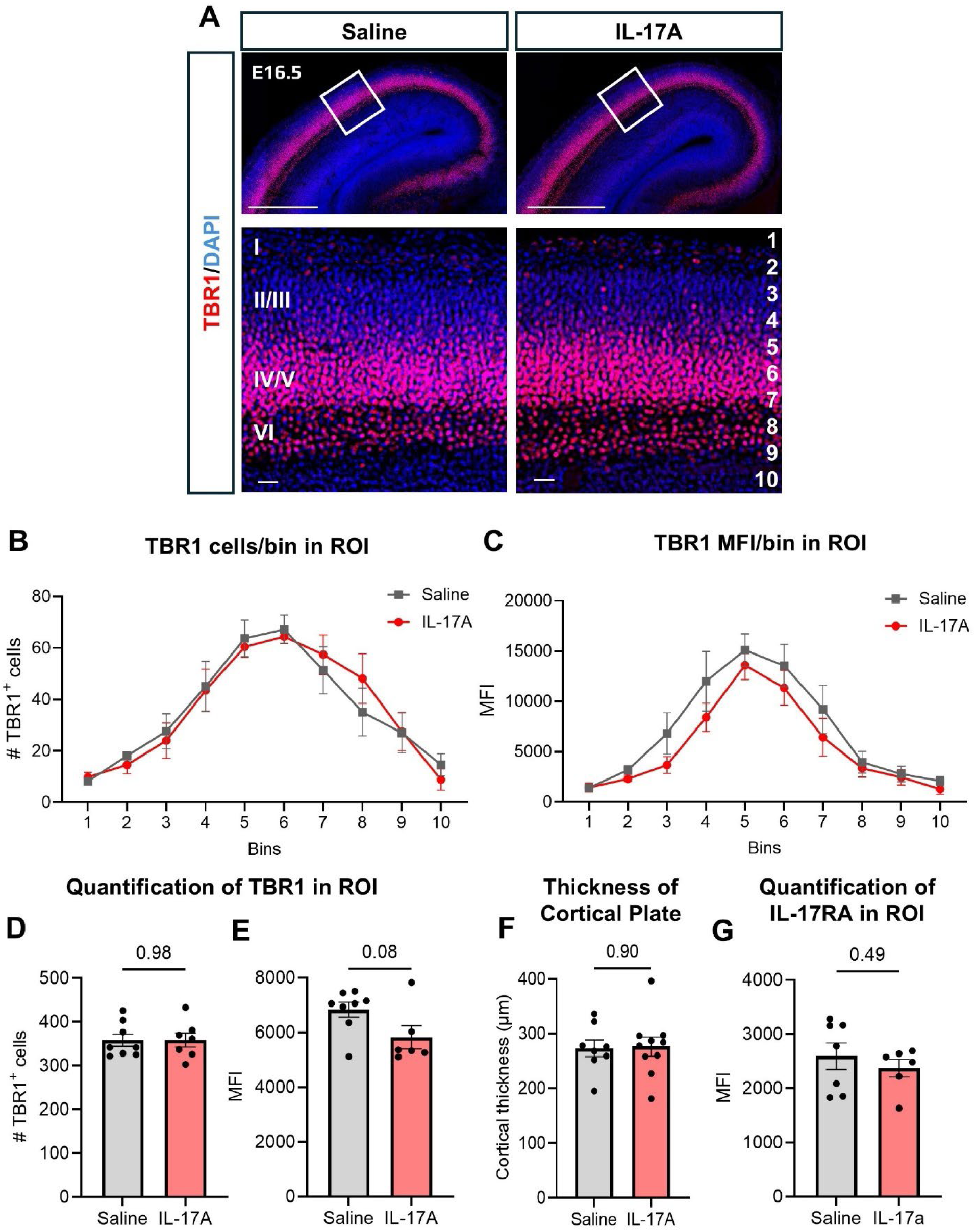
Gestational IL-17A administration did not alter the structure of the fetal primary somatosensory cortex. **(A)** Representative GD16.5 fetal brain section immunostained for TBR1 (red) and counterstained with DAPI (blue). Images were taken from the right-hemisphere somatosensory cortex within a 300 × 300 µm² ROI. Top scale bars = 500 µm; bottom scale bars = 20 µm. Numbers at the left indicate cortical layers, numbers at the right indicate bin divisions. **(B)** IL-17A administration did not alter the distribution of TBR1⁺ cells across the cortical plate. **(C)** IL-17A administration did not alter the mean fluorescence intensity of TBR1 across the cortical plate. **(D)** IL-17A administration did not alter the total number of TBR1⁺ cells in the ROI. **(E)** IL-17A administration did not alter the mean fluorescence intensity of TBR1 in the ROI. **(F)** IL-17A administration did not alter the total cortical plate thickness. **(G)** Administration of IL-17A did not alter the mean fluorescence intensity of IL-17RA in the cortical plate. Data are means ± SEM. Saline: n = 7-8; IL-17A: n = 6-10.

Although we confirmed that rIL-17A can stimulate brain microglia-like cells *in vitro* **(Supp. Fig. S1)**, it was possible that the developing GF fetal brain may not have the necessary machinery to respond to IL-17A *in vivo*. To test this, we measured IL-17A receptor subunit A (IL-17RA), which complexes with IL-17RC, forming the heterodimer that binds IL-17A. Previous studies have demonstrated that transcripts for IL-17RA, but not IL-17RC, increase in the fetal brain in response to poly I:C-induced MIA [22, 23]. In GF fetal brains, we observed no changes in IL-17RA protein—assessed via fluorescent immunohistochemistry—in response to maternal rIL-17A administration **(Fig. 3G)**. Notably, IL-17RA was detectable in all GF brains, with positive expression throughout the developing cortex, indicating that GF E16.5 fetuses likely have the capacity to respond to IL-17A.

To more extensively characterize cortical lamination in E16.5 GF fetal brains, we utilized spatial light interference microscopy (SLIM). SLIM is a label-free, quantitative phase imaging modality that renders optical pathlength (topographic) maps sensitive to tissue cytoarchitecture, enabling morphological assessment without the use of exogenous stains [69]. To illustrate correspondence with conventional histology, we paired SLIM images with matched H&E bright-field images of the same E16.5 sections, as shown in Supp. Fig S2. Sections of the whole fetal brain at 5X **(Supp. Fig. S2A,B)** demonstrate detailed cytoarchitectural organization with clear layer delineation, showing that SLIM accurately preserves and enhances the structural features visible in conventional H&E images. In the ROI-scale views at 10X and 40X **(Supp. Fig. S2C-F)**. Across these scales, SLIM recapitulated the laminar organization evident in H&E-stained slices, supporting its use for topology-based quantification of fetal cortical cytoarchitecture.

To evaluate the morphological consequences of maternal IL-17A administration, cellular density was quantified in the E16.5 primary somatosensory cortex. Using SLIM, images of the ROI were captured at 10X **(Fig. 4A)** and uploaded to a trained Biodock project for automated event segmentation. Total events (i.e., individual cell bodies) and event density (events/mm²) were calculated for each cortical layer (I–VI). Across each of the cortical layers (I, II/III, IV/V, VI), overall events did not differ between fetuses of control and IL-17A–treated dams **(Fig. 4B)**. As an orthogonal validation, normalized DAPI fluorescence intensity (i.e., nuclei abundance) measured from adjacent sections using the same automated event quantification method showed no group differences between control and IL-17A–treated offspring **(Fig. 4C)**. An example Biodock-generated map of individually quantified cell “events” is shown in **Fig. 4D**. Together, these analyses show that label-free phase morphology captures embryonic cortical lamination comparable to traditional H&E histology approaches, the first time this has been demonstrated. Moreover, this approach further confirms that IL-17A administration in pregnant GF dams fails to produce dysplasia-like signatures in the E16.5 somatosensory cortex.

**Figure 4.**
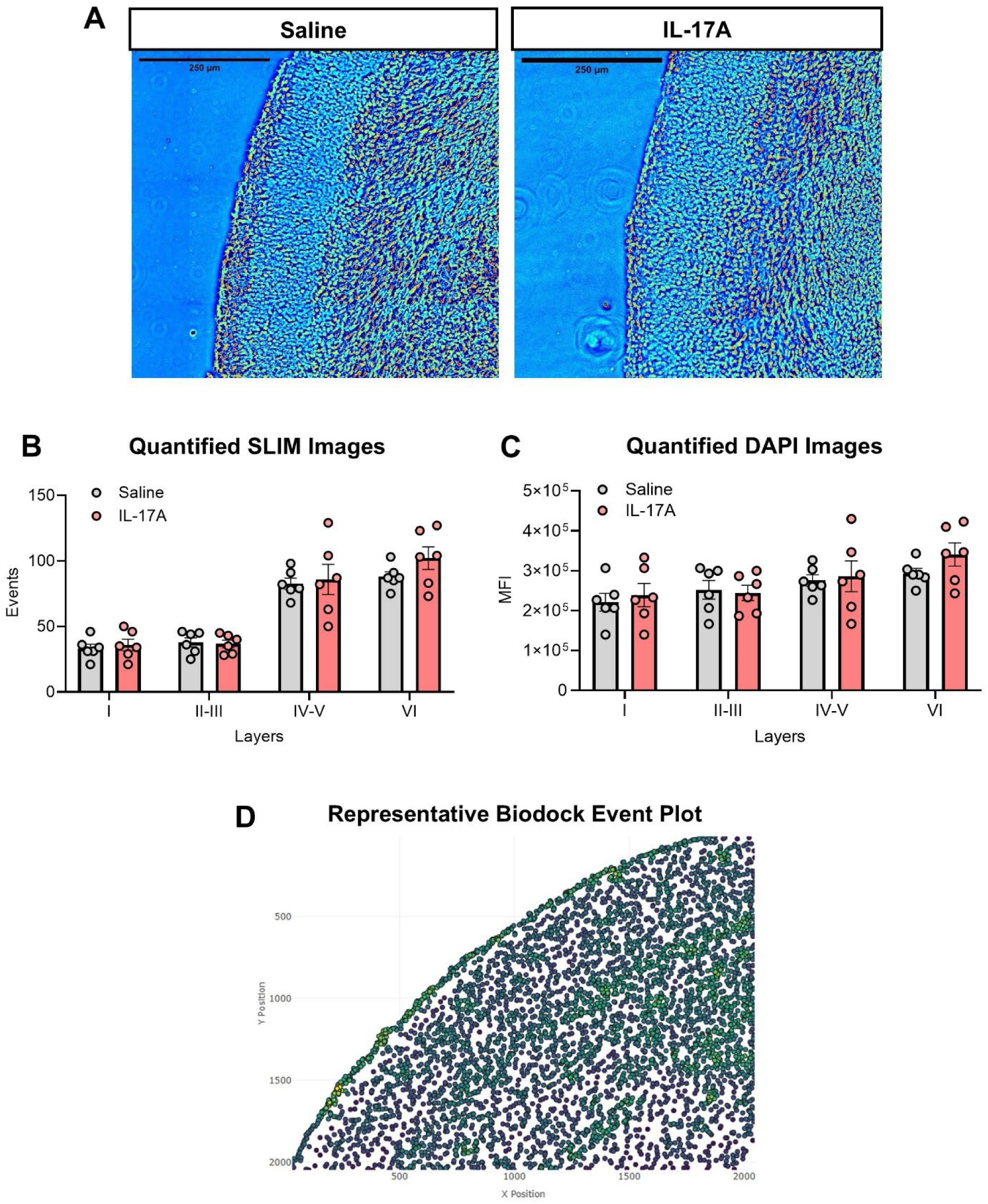
Gestational IL-17A administration did not alter cell density distribution across cortical layers of the fetal primary somatosensory cortex. **(A)** Representative GD16.5 fetal brain images from saline- and IL-17A-treated dams, showing the right-hemisphere somatosensory cortex within a 300 × 300 µm² ROI. Scale bar = 250 µm, magnification 10X. **(B)** IL-17A administration did not alter the total number of events per cortical layer. **(C)** IL-17A administration did not alter cellular density across cortical layers. **(D)** Biodock-generated scatterplot showing event counts derived from SLIM image data, where blue-to-yellow color represents increasing density. Data are means ± SEM. Saline: n = 6; IL-17A: n = 6.

## 4.0 Discussion

Using repeated administration of rIL-17A in GF mice to mimic late-gestation maternal inflammation, we demonstrate that maternally derived IL-17A is insufficient for disrupting fetal cortical lamination in the absence of microbes. In comparison to MIA studies demonstrating abnormalities in the fetal somatosensory cortex following IL-17A-dominant signaling, our findings indicate that microbes (and the mature immune profiles that come with them) may be required for this type of neurodevelopmental disruption. To our knowledge, this is also the first time that spatial light interference microscopy (SLIM) has been applied to the embryonic mouse brain, opening avenues for future use of this technique on non-labeled fetal tissue.

It should be noted that our findings do not undermine the potential role of IL-17A, in conjunction with other signaling molecules, as a co-mediator of NDD-like pathogenesis during maternal inflammatory conditions. Instead, our study further emphasizes the importance of the microbiome during prenatal development and highlights the merit of distinguishing microbial-derived signals from immune-derived signals when investigating NDD etiologies. IL-17A has been identified as a key factor that can induce NDD-like phenotypes in murine models, including disrupted cortical lamination and behavioral changes *in vivo*, and similar defects after direct fetal brain exposure [22, 74, 75]. In this study, we administered 1µg of rIL-17A for six consecutive days of gestation with the intention of mimicking mid-gestation maternal viral infection. Circulating IL-17A in GF mice reached levels comparable to those induced by well-established MIA models utilizing viral mimic poly I:C [22]. Thus, if circulating maternal IL-17A alone, independent of microbes, is sufficient to induce fetal cortical abnormalities similar to those previously reported [22, 23, 39], we would expect to observe them here. The fact that cortical architecture remains intact, at least in the primary somatosensory cortex, indicates that IL-17A alone is insufficient for overtly disrupting lamination patterns. However, it is important to note that we were unable to confirm whether circulating IL-17A levels in the fetus recapitulated those in the pregnant dam. Yet, this is often not reported in poly I:C MIA models either. What has been reported is an increase in IL-17 receptor transcripts in the fetal brain [22], indicating ligand-induced receptor upregulation. Here, we do not observe an increase in fetal brain IL-17RA protein as measured by immunohistochemistry. Based on our *in vitro* BV-2 stimulation experiment, we expect rIL-17A to be biologically active and capable of signaling in the brain. Thus, these findings could indicate that microbes may be required for rIL-17A to induce receptor upregulation. Alternatively, it could indicate that rIL-17A administered i.p. in pregnant dams is not reaching the fetal brain. To truly determine whether maternal IL-17A directly reaches the fetal brain, in a GF model like ours and in MIA models, future studies are needed to rigorously trace maternally derived cytokines across the maternal-fetal interface and into the fetal compartment. It is also possible that IL-17A signals locally at the placenta, propagating further IL-17 release within the fetal compartment [32, 76]. Timing of IL-17A signaling (E10.5 to 15.5 in our model versus E12.5-14.5 in others [22, 23]) is also likely to play a role. Moreover, an intracerebroventricular administration of rIL-17A into the fetal brain (as reported previously in specific pathogen free [SPF] mice [22]) may be needed to definitively determine whether IL-17A is capable of directly disrupting neurodevelopment in GF mice.

The role of the gastrointestinal microbiome in MIA has been highlighted by recent studies demonstrating that microbiome composition is a modulator of the inflammatory response and can predict MIA outcomes [39, 74, 77]. Furthermore, the presence or absence of specific bacteria, such as segmented filamentous bacteria, determines whether MIA induces maternal TH17 cell expansion and IL-17A production, thereby critically regulating MIA outcomes [43]. Consistently, other work has shown that specific commensal taxa, notably SFB, drive the differentiation of small-intestinal TH17 cells, which are the principal source of IL-17A [74]. Consistent with this, our findings in GF dams align with previous work showing that poly(I:C)-induced MIA does not disrupt cortical development in the absence of SFB [39], as this microbial priming is required for maternal TH17 expansion and IL-17A responses. In our model, IL-17A administration was similarly insufficient to disrupt fetal cortical lamination in GF mice, underscoring the potential role of the microbiome in enabling IL-17A-induced effects. Moreover, it has been shown that transfer of gut microbiota from MIA donors is sufficient to reproduce some MIA-associated neuropathological and inflammatory features in offspring [78]. While the mechanisms underlying microbial regulation of NDD pathogenesis are complex and still unclear, recent research suggests that microbial metabolites (e.g., TMAO, 5-AV, IP, and HIP) can directly promote axon outgrowth in fetal thalamic explants and rescue axonogenesis *in vivo*, thereby impacting brain development [34]. In addition to supporting fetal brain development, microbial metabolites similarly regulate placental development, providing a parallel route through which the maternal microbiome can shape fetal developmental outcomes [34, 35]. Taken together, these observations indicate that the absence of a maternal microbiome removes the upstream signals necessary for IL-17A-mediated neurodevelopmental effects, an intriguing potential explanation for why exogenous rIL-17A alone failed to alter cortical lamination.

There are several key differences that should be noted between our study and most murine models of MIA. In studies of maternal infection with live pathogens, dams show suppressed gestational weight gain, and litter characteristics are often altered [67, 79, 80]. This weight suppression can reflect inflammation-mediated sickness behavior and the energetic cost of a systemic immune response. In previous work from our lab using influenza A infection, attenuation in maternal weight gain accompanied increases in systemic inflammation [53, 66, 67]. In contrast, our IL-17A administration paradigm tests the effect of IL-17A alone, without inducing the broader immune response to infection, which explains the absence of maternal weight loss and altered litter outcomes. Consistent with the absence of a full anti-pathogen response, we observed no evidence of systemic inflammation as neither spleen weight nor colon length differed from controls.

Regarding placenta structure, it is important to highlight that GF mice lack the microbial derived signaling compounds that promote placental growth and vascularization [35]. In this study, we did not directly compare outcomes with traditional SPF laboratory mice; thus, we cannot say whether differences between GF and SPF placental development may contribute to our findings. The histological and morphometric approaches used in this study are well-established methods for detecting placental malformations and pathological alterations [48–51, 53]. Furthermore, previous studies have shown that MIA can cause alterations in the placenta and still fail to induce changes in fetal neurological development [53, 59]. Another aspect to consider regarding placental morphological responses is the possibility of sex-specific effects, as other studies have reported that maternal inflammation induces a male-biased placental response, with male placentae showing stronger up-regulation of cytokines such as IL-6 and IL-1β compared to females [81]. In addition, human studies have shown sex-specific epigenetic alterations in intrauterine growth restriction, suggesting distinct mechanisms of placental vulnerability according to fetal sex [82]. In our study, we chose to assess only one placenta per litter to avoid sample size inflation, but this precluded assessment of sex-specific effects, which therefore cannot be ruled out. Future studies powered to distinguish sex effects are required to sufficiently parse this out in both the placenta and fetal brain.

Evaluation of cortical structure was conducted using two methods: TBR1⁺ cell distribution and cellular density across layers quantified by Spatial Light Interference Microscopy. In both analyses, maternal IL-17A had no effect. TBR1⁺ immunostaining has been widely used to assess cortical layering and excitatory neuron positioning, providing a common readout for cortical lamination and structural integrity [22, 39, 67]. While SATB2 (a marker for upper-layer excitatory neurons) is often paired with TBR1 (deep-layer excitatory neurons) to reveal MIA-associated lamination effects, persistent changes in TBR1 have been reported in both poly I:C- and viral-induced MIA models. Both Choi *et al.* (2016) and Otero *et al.* (2025) report bin-specific alterations in TBR1 distribution [22, 67]; in a follow-up study by Choi’s group, scattered areas of “cortical patches” defined by the absence of TBR1 are also reported [23]. In our paradigm, TBR1 distribution was unchanged, indicating that maternal IL-17A exposure in the absence of microbes does not produce overt deficiencies in deep-layer laminar cytoarchitecture. While it is possible that rIL-17A administration may still impact upper-layer neurons in GF mice, we believe that our additional complementary measures of cortical cytoarchitecture indicate that neuronal lamination patterns across all layers are mostly conserved.

We used SLIM, a quantitative phase imaging technique, to provide wide-field mosaics of whole sections and to delineate discrete cells within each cortical layer [69]. This label-free, high-resolution imaging technique is particularly well-suited to detect disruptions such as focal patches or cytoarchitectural abnormalities commonly associated with NDDs, which would manifest as localized changes in cell density across cortical layers [69, 83]. In this study, SLIM used to identify high contrast individual cellular events that we used as a proxy for cortical cell density. This constitutes a novel application of SLIM to embryonic brain histology, where it serves as a complementary label-free quantitative approach to conventional immunohistochemistry. Given the absence of focal cortical disorganization as revealed by SLIM, we surmise that any layer-specific alterations detectable with neuron-specific markers would be minute, if present at GD 16.5.

Overall, our results indicate that, without microbial priming, IL-17A alone is insufficient for disrupting placental and fetal brain development, highlighting the maternal microbiome as the critical determinant to target and control in MIA models. However, we also recognize that the simplicity of our study design limits the extent of the conclusions that can be made. For example, while overt structural defects may be absent in the cortical region of interest here, it is possible that cytoarchitectural abnormalities exist in other brain areas or that behavioral abnormalities may yet manifest postnatally. Further studies are needed to directly test these possibilities. Still, we believe this work sets the foundation for further rigorous examinations designed to parse out whether specific developmental outcomes are dependent upon immune or microbial signaling, or the combination of both. Overall, our study adds to the growing literature indicating that the maternal microbiome likely acts in concert with the maternal inflammatory response when impacting fetal development.

## 5.0 Declarations

### 5.3 Data and Code Availability

All data supporting the findings of this study are available within the paper and its Supplementary Information. No code was utilized.

### 5.4 Authors’ Contributions (CRediT author statement) and Acknowledgements

**Izan Chalen:** Methodology, Formal analysis, Investigation, Writing - Original Draft. **Selena Wang:** Methodology, Investigation. **Rafael J. Gonzalez-Ricon:** Investigation. **Ashley M. Otero:** Investigation. **Fernando J. Rigal:** Investigation. **Alexander Byrne:** Investigation. **Shakirat Adetunji:** Investigation. **Catherine Best-Popescu:** Resources, Supervision. **Adrienne M. Antonson:** Conceptualization, Writing - Review & Editing, Project administration, Funding acquisition. We thank Dr. Robert McCusker for his kind donation of mouse BV-2 cells.

### 5.5 Registration details

NOT APPLICABLE (not a clinical trial). **Ethics and Consent to Participate declarations:** not applicable. **Consent to Publish declaration:** not applicable.

---

**Supplementary Figure S1.**
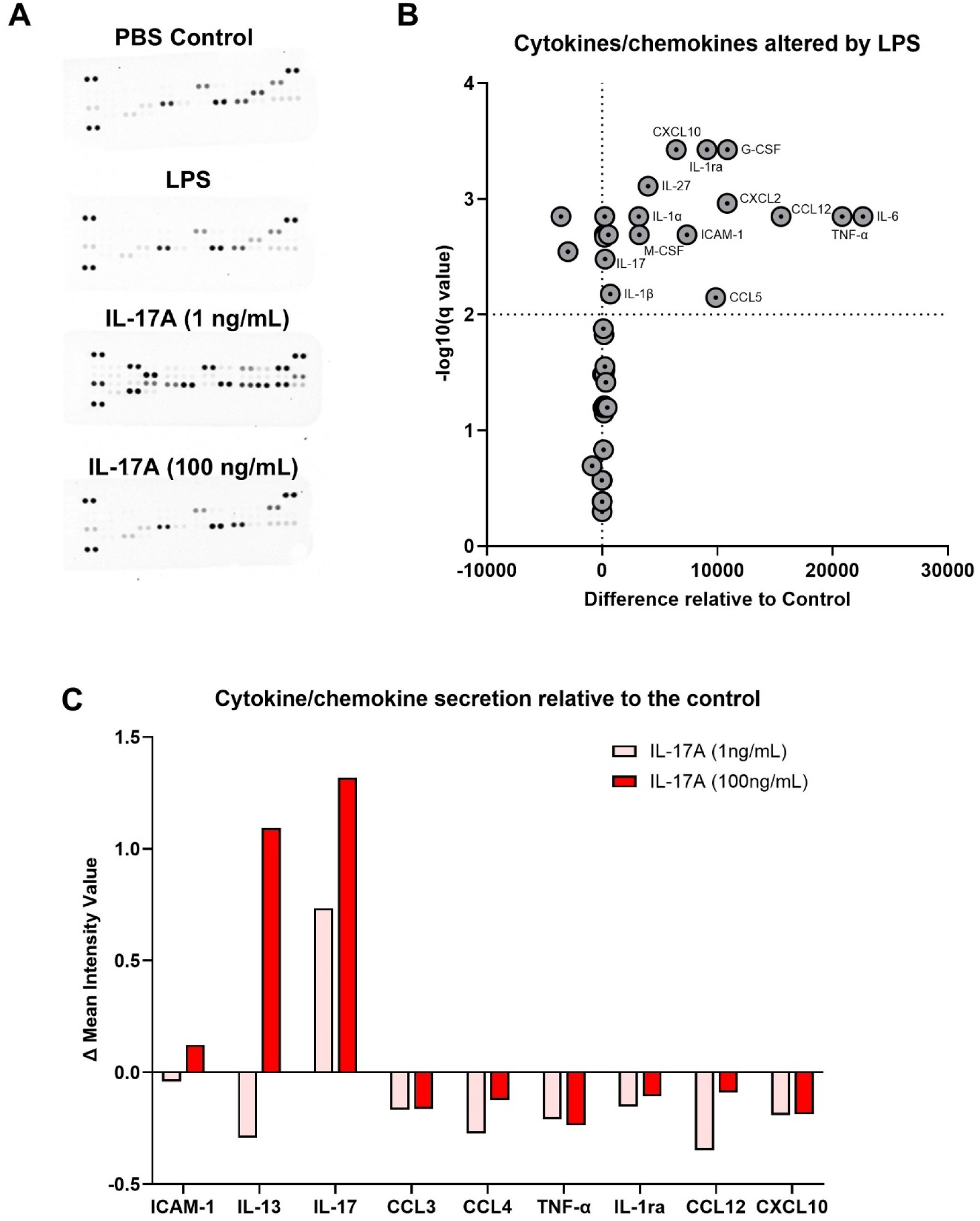
Recombinant mouse IL-17A is bioactive in BV-2 microglial cells *in vitro*. **(A)** Proteome profiler membranes showing cytokine and chemokine profiles from PBS-treated (negative control), LPS-treated (positive control), and recombinant IL-17A–treated BV-2 cells at 1 ng/mL (low dose) and 100 ng/mL (high dose). **(B)** Volcano plot showing the protein response to LPS stimulation relative to PBS control. **(C)** Bar graph showing relative expression levels of selected cytokines in BV-2 cells stimulated with low- or high-dose IL-17A (1 or 100 ng/mL) compared with PBS control.

**Supplementary Figure S2.**
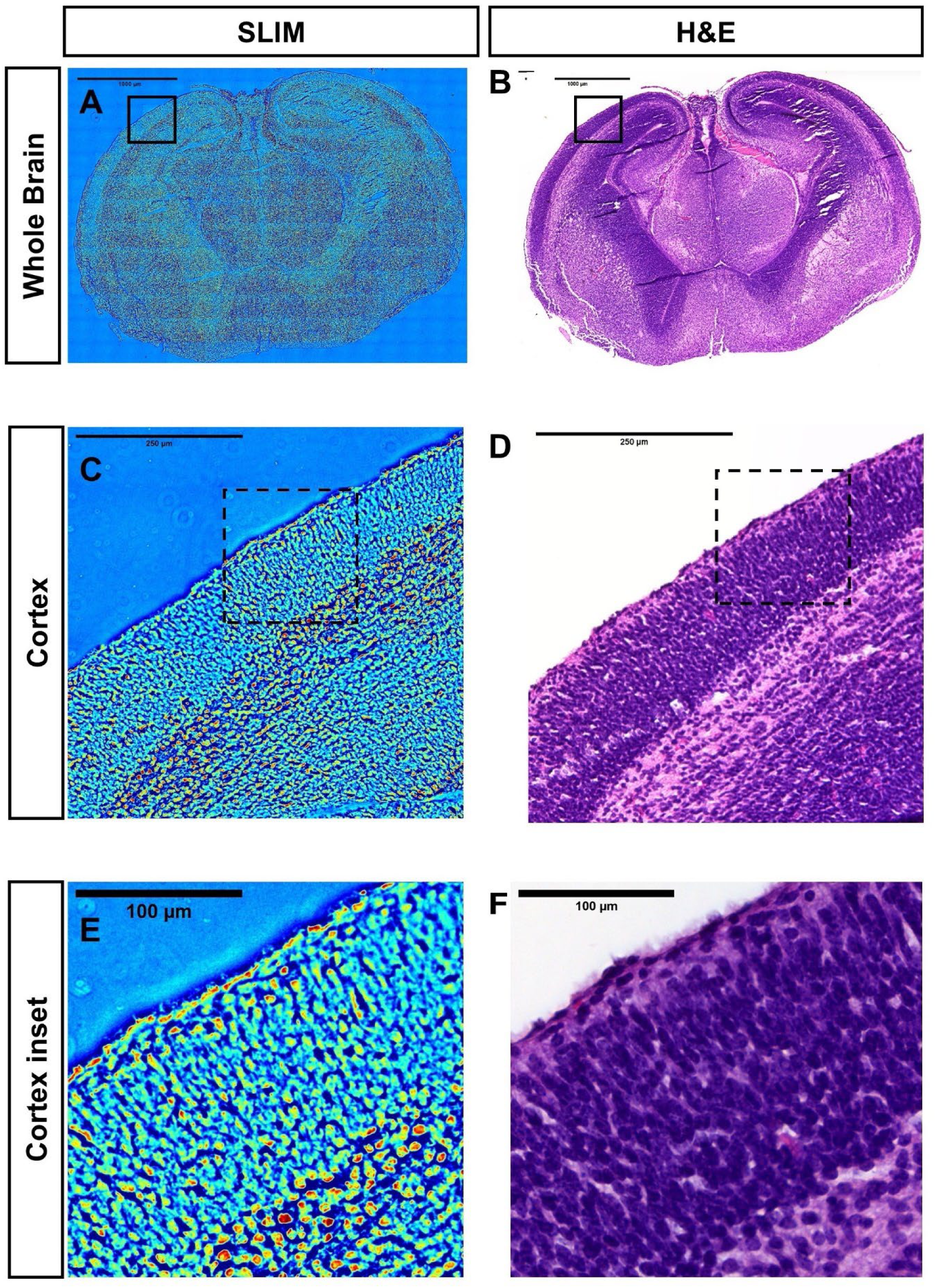
SLIM fetal brain images compared to traditional H&E histology. **(A)** Whole coronal section of a GD16.5 fetal brain imaged using Spatial Light Interference Microscopy (SLIM). The region of interest (ROI) is indicated by a rectangle. **(B)** Corresponding H&E-stained brightfield image of the same coronal section shown in (A). **(C)** ROI from a GD16.5 fetal brain coronal section imaged under SLIM. **(D)** H&E-stained brightfield image corresponding to the ROI shown in (C). **(E)** Magnified SLIM image of the cortical region indicated in (C). **(F)** Corresponding H&E-stained magnified brightfield image of the same cortical region shown in (E).

